# A mannitol/sorbitol receptor stimulates dietary intake in *Tribolium castaneum*

**DOI:** 10.1101/157347

**Authors:** Tomoyuki Takada, Ryoichi Sato, Shingo Kikuta

## Abstract

Perception of chemical stimuli by insects aids in accepting or rejecting food. Gustatory receptors (Grs) regulating external signals in chemosensory organs have been found in many insects. *Tribolium castaneum*, a major pest of stored products, possesses over 200 *Gr* genes. An expanded repertoire of *Gr* genes appears to be required for diet recognition in generalist feeders; however, it remains unclear whether *T. castaneum* recognizes a suite of chemicals common to many products or whether it is attracted to specific chemicals, and whether its Grs are involved in its feeding behavior. It is difficult to determine the food preference of *T. castaneum* based on its dietary intake due to a lack of appropriate methodology. This study established a novel dietary intake estimation method using gypsum, designated the TribUTE (*Tribolium* Urges To Eat) assay. *T. castaneum* adults were fed gypsum block without added organic compounds. Sugar preference was determined by adding sweeteners and measuring the amount of gypsum in the excreta. Mannitol was the strongest attractant of *T. castaneum* dietary intake; in addition, TcGr20 was responsible for mannitol and sorbitol responses in *Xenopus* oocyte expression, but did not respond to any other non-volatile compounds tested. The EC_50_ values of TcGr20 for mannitol and sorbitol were 72.6 mM and 90.6 mM, respectively, suggesting that TcGr20 is a feasible receptor for the recognition of mannitol in lower concentrations. *TcGr20* was expressed in the antennae, where the perception of mannitol would occur. We examined whether *TcGr20* was involved in mannitol recognition using RNAi and the TribUTE assay. The amounts of excreta in *TcGr20* dsRNA-injected adults decreased significantly despite the presence of mannitol, compared to that of the control adults. Taken together, our results suggest that *T. castaneum* adults recognized mannitol/sorbitol using TcGr20 receptors, thereby facilitating their dietary intake.

**Abbreviation:**

ORF: open reading frame
RT: reverse transcription
RNAi: RNA interference
dsRNA: double strand RNA
cRNA: capped RNA
MBS: modified Barth’s saline
CBB: Coomassie brilliant blue
Gr: gustatory receptor

## Introduction

Feeding behavior in insects is comprised of several processes for recognizing chemical compounds, tasting, continuous feeding, and digestion [1]. Food-acceptance or food-rejection actions in insects are determined by non-volatile compounds such as carbohydrates and caffeine contained in host plants [2, 3]. Chemical compounds stimulate the gustatory receptors (Grs) located in external sensory organs. Stimulated Grs transmit electrochemical signals through sensory neurons to the sub-esophageal ganglion and brain, thereby regulating the feeding behaviors of insects [4, 5]. Feeding behaviors differ across diverse insect species [6]. For example, larval growth of the specialist-feeding silkworm *Bombyx mori* depends solely on mulberry leaves [7]. *B. mori* perceives some gustatory stimulants such as sucrose, inositol, morin and (β-sitosterol contained in the mulberry leaves using maxillary palps [8]. *Myo*-inositol, an indispensable nutrient and a feeding behavior prolonging factor in *B. mori* larvae, was recognized via BmGr10 expression in the sensory organs [9]. Therefore, Grs in specialist feeders play a key role in detecting specific compounds. Meanwhile, a generalist feeder, *Helicoverpa armigera*, utilizes a wider range of host plants. Given that generalists feed on various plants and plant products, many types of non-volatile chemical compounds contained in foods may be sensed by various Grs-expressing sensory organs [10,11].

*Tribolium castaneum*, a major pest of grains, cereals, pasta, chocolates, and nuts [12,13], possesses 207 Grs genes available as gene models based on genome analysis [14]. Tissue-specific expression analyses have shown that 34 *Grs* genes in the antennae associated with the gustatory perception, with many more types of Grs expressed in antennae than in other insect species [15]. This expanded Gr family likely plays a functional role for host selection. One pressing question pertains to whether *T. castaneum* recognizes a suite of chemicals common to many products, or whether it is attracted to specific chemicals. To answer this question, a dietary intake evaluation of *T. castaneum* is required using an artificial diet composed of as few compounds as possible. Dietary intake of *T. castaneum* has been previously examined using dried flour, but this is insufficient for evaluating preferences since the organic compounds in flour cannot be completely separated [16,17]. Therefore, the establishment of a dietary intake assay in an organic-free state is required. One commonly-used method involves the measurement of swallowed liquid food composed of sugar and water, such as in the CAFE (Capillary Feeder) assay [18]. However, this method is only applicable for sucking or licking insects. Because *T. castaneum* prefers dry products with water content less than 12% [17,19], CAFE cannot be applied for investigations into its dietary preference. Little is known about the dietary intake of *T. castaneum* due to the lack of technical methods using dry compounds. This study developed a novel dietary intake estimation method using gypsum block without added organics. Since the gypsum eaten by *T. castaneum* adults is eventually excreted without digestion as waste, the measurement of excreta allows for the quantification of dietary intake in *T. castaneum*.

Additionally, sugar preference can be determined by adding sweeteners to act as a stimulant for feeding behavior. Sweeteners would act as ligands to stimulate Grs; however, to date, only few Grs have been identified as ligand-stimulated in the ectopic system in generalist feeders, including *T. castaneum*. HarmGr4, found in *H. armigera*, responded to fructose in the *Xenopus* oocyte expression system [11]. Furthermore, the Gr43-like clade in Gr family including HarmGr4, have been found in various insects such as the Diptera and Lepidoptera, where they act as receptors of fructose and *myo*-inositol [9, 20]. Since the Gr43-like genes are also represented in *T. castaneum*, the candidate genes may be stimulated by any sugars/sugar alcohols. Here, we predicted that sweeteners deduced in the dietary intake assay were potentially ligand stimulators of Gr43-like genes in *T. castaneum*. This study demonstrates ligands of TcGr20 belonging in the Gr43-like gene family using an exogenous expression system with a two-voltage clamp assay. We also demonstrate the *in vivo* effect of the combination of RNAi and the dietary intake assay.

## Materials and methods

### Insects

Red flour beetles (*Tribolium castaneum* Herbst.) were obtained from Sumica Technoservice Co. (Hyogo, Japan) and reared on whole-wheat flour (Pioneer-kikaku, Kanagawa, Japan) and yeast (Saf-instant®, Lesaffre, Marcq-en-Baroeul, France). They were maintained at 29 ± 1°C under a 16 L:8 D cycle.

### Total RNA preparation and cDNA synthesis

Total RNA was isolated from the antennae and proboscis of 20 adults using ISOGEN II (NIPPON GENE, Tokyo, Japan) following the manufacturer’s instructions. cDNA was synthesized from total RNA by ReverTra Ace® (TOYOBO, Osaka, Japan) using Oligo-dT primers. Primer sequences are shown in S1 Table. The cDNA transcription reaction was performed at 42°C for 90 min to produce cDNA and 99°C for 5 min to denature the enzymes. Open reading frames (ORF) of *TcGr20, TcGr21, TcGr27 and TcGr28* were amplified from cDNA by PCR using a high-fidelity DNA polymerase, PrimeSTAR® HS (TaKaRa Bio, Shiga, Japan) with specific primers containing restriction enzyme sites at 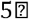 and 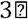 ends of ORF (S1 Table). Kozak sequences were inserted between restriction sites and primer binding sites to enhance translational efficiency. PCR reactions were performed as follows: 35 cycles of denaturation at 98°C for 30 s, annealing at 55°C for 15 s, and extension at 72°C for 70 s. *TcGr20, TcGr21, TcGr27* and *TcGr28* were directly subcloned into expression vector pT7XbG2 (DDBJ accession number, AB255037) and the sequence analyses were performed by Eurofins Genomics (Tokyo, Japan) to confirm the correctness of the construct. Sequence data were analyzed using FinchTV sequence scanner software.

### Quantitative RT-PCR

Total RNA was isolated from male and female adults (n = 20) with ISOGEN II (NIPPON GENE, Tokyo, Japan) following the manufacturer’s instructions. Gene expression levels were examined in various tissues: antennae, heads (without antennae but including mouthparts), thorax, abdomen and legs. First-strand cDNA was synthesized using a PrimeScript™ RT reagent Kit (TaKaRa Bio) following the manufacturer’s instructions. The quality and concentration of synthesized cDNA were measured using a NanoPhotometer® NP80 (Implen, München, Germany). Quantitative RT-PCR was performed using SYBR® Premix *Ex Taq™* II (Tli RNaseH Plus, TaKaRa Bio) with a StepOnePlus™ (Thermo Fisher Scientific, Carlsbad, CA) under the following conditions: a holding cycle at 95°C for 10 min, followed by 40°C cycling stage of 95°C for 15 s, 60°C for 1 min, and melting curve stages were carried out at 95°C for 15 s and 60°C for 1 min to confirm the presence of nonspecific PCR reactions. Relative expression levels were calculated using *ΔΔCt*. The *TcGr20* expression levels in tissues were normalized using the expression levels of *T. castaneum* ribosomal protein S3 (*RpS3*, NCBI accession: NM_001172392.1). *RpS3* was used as a housekeeping gene due to the expression stability through the stage from larvae to adult [21, 22]. Primer sequences are shown in S2 Table.

### Capped RNA synthesis and two-electrode voltage clamp electrophysiology

Procedures followed those described in a previous study [9]. Briefly, capped-RNA (cRNA) were synthesized using mMESSAGE mMACHINE® T7 kit (Thermo Fisher Scientific) according to the manufacturer’s instructions, and were kept at 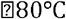 until use. The cRNA injected oocytes were incubated in modified Barth's saline (MBS) buffer supplemented with 10 mg/mL penicillin and streptomycin for 3 days at 20°C [23]. Water-injected oocytes, which had endogenous receptor activities alone, were used as negative controls. Whole-cell current was recorded with two-electrode voltage clamp in a perfusion system using Ringer’s solution [9]. Current was amplified with an OC-725C amplifier (Warner Instruments, Hamden, CT, USA) at a holding potential of 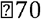 mV, low-pass filtered at 50 Hz, and digitized at 1 kHz. Data was acquired by software pCLAMP™ 10 (Molecular Devices, Sunnyvale, CA). Dose-dependent response curves were fitted to a hill slope curve using Prism 6 software (GraphPad, San Diego, CA).

### Evaluation of dietary intake

We developed a novel feeding application for *T. castaneum* adults using gypsum containing a near-zero amount of organic compounds instead of artificial diets and wheat, termed a TribUTE (*<u>Trib</u>olium* <u>U</u>rges <u>T</u>o <u>E</u>at) assay. The artificial gypsum diets were comprised of dry powder and water mixed in a ratio of 1.3:1 (w/w) containing 200 mM sugars or sugar alcohol solutions. The mixed gypsum was allowed to solidify completely at 65°C for 48 h. *T. castaneum* adults were starved for one week in the absence of foods and held in cages at 25°C. Each gypsum block of approximately 5 mm was provided to individual adult beetles in a 24-well microplate. They were kept for 48 h at 25°C. The eaten gypsum by *T. castaneum* adult was eventually excreted without digestion as a waste, permitting the measurement of the amount of gypsum. The excrements of the artificial gypsum diet were collected in 200 μL microtubes by using a thinning silicon wire under a stereoscopic microscope. The excreta were dissolved with 50 μL deionized water to remove sugars or sugar alcohols, and then the precipitate of gypsum was dried thoroughly at 65°C for 24 h. Excreta were weighed by microbalance (AT201, Mettler-Toledo, OH).

### RNA interference

To silence the gene expression of *TcGr20* in adults, we synthesized double-strand (dsRNA) *in vitro* using MEGAscript® T7 RNAi kit (Thermo Fisher Scientific) according to the manufacturer’s instructions. Primers sequences are shown in S3 Table. The dsRNA at 1 μg/μL was kept at -80°C until use. A 150-200 nL volume of dsRNA was injected between the internode of head and thorax in adult beetles using a capillary needle, Nanoject II (Drummond Scientific Company, Broomall, PA) on a cooling block at -10°C. Emerald luciferase (*Eluc*, TOYOBO, Osaka, Japan) was used as a negative control. The similar sequences of *Eluc* were searched in NCBI *Tribolium* genome (ID:216). Since the over 22-bp identical sequences of *Eluc* and *TcGr20* could potentially function as false targets, these sequences were trimmed from the dsRNA regions. The dsRNA-injected adults were kept at 25°C. Adult beetles were used for the TribUTE assays and quantitative RT-PCR at 48 h after injections.

## Results

### Identification of feeding promoting stimulants in *T. castaneum*

We found that *T. castaneum* adults consumed gypsum block in the absence of organic compounds (Fig 1A). To discriminate gypsum excreta from conventional excreta derived from wheat flour, gypsum block stained with Coomassie Brilliant Blue (CBB) R-250 (Wako, Osaka, Japan) was given to adult beetles (Fig 1A). We visualized the digestive tract from the foregut to the anus under a microscope, and found that the tract was partially stained by CBB (Fig 1B), while the tract was not stained using gypsum without CBB (Fig 1B’). The excreta of the stained gypsum were also observed (Fig 1C). Together, these findings showed that *T. castaneum* adults recognized and fed on gypsum, and excreted it as a waste. We then attempted to identify feeding-promoting stimulants for *T. castaneum*. We produced gypsum blocks containing mono-, di-, and tri-saccharides or sugar alcohols as candidates. As the amount of excreta with these additives was higher than the amount produced from sugar-free gypsum, we assumed that *T. castaneum* recognized sugars and sugar alcohols as attractants. *T. castaneum* actively fed on gypsum with added sugars/sugar alcohols, and excreted the gypsum as a waste. The amount of excreted sweetened gypsum was significantly greater than gypsum without sweetener (Fig 1D). Additionally, gypsum with 200 mM galactose and sucrose yielded a negligible and non-significant difference compared to gypsum without sweeteners (Fig 1D). The amount of excreta was markedly greater in the presence of 200 mM mannitol, demonstrating that feeding behavior was stimulated based on the sweetener itself. These findings indicate that mannitol acts as a strong attractant for *T. castaneum*.

**Fig 1.**
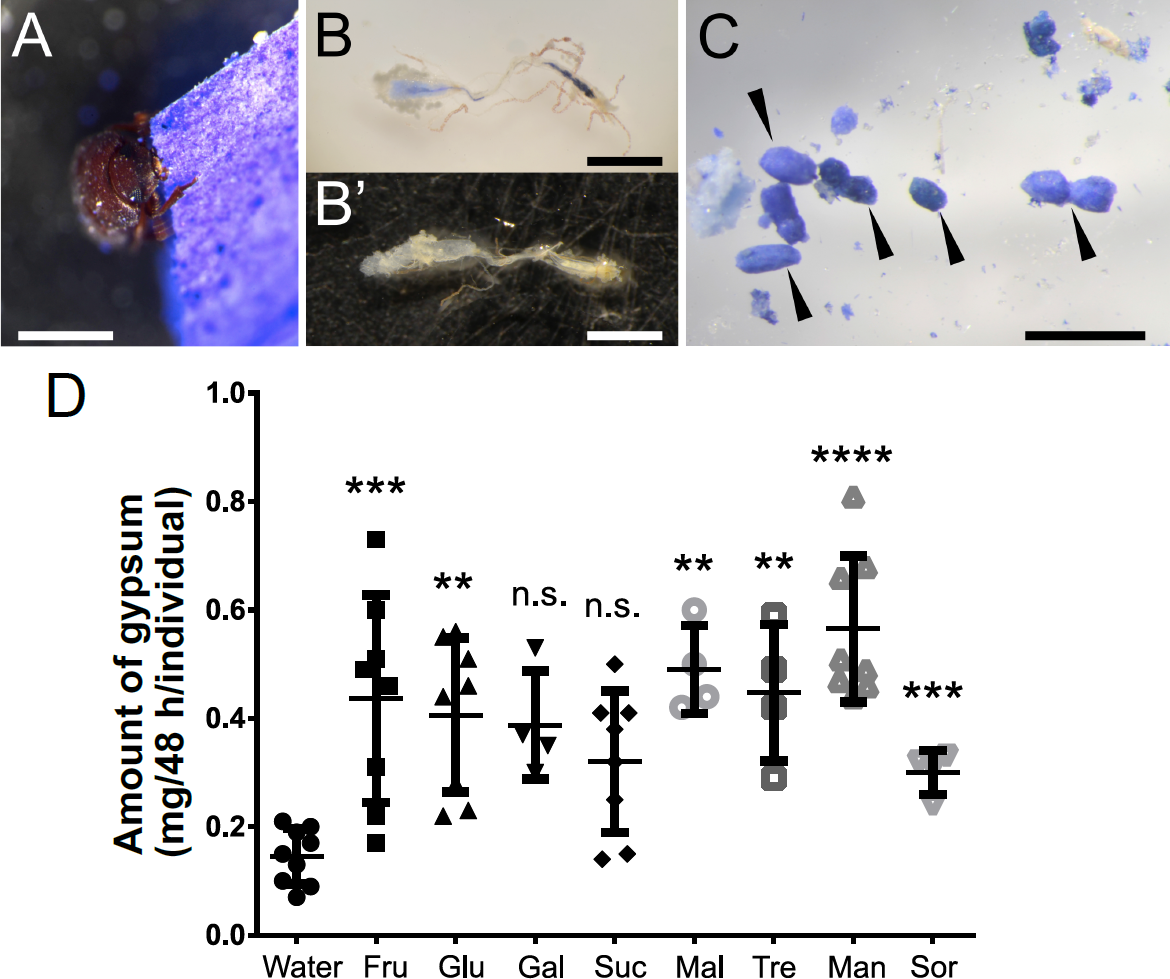
Establishment of feeding assay in *T. castaneum* adults using gypsum block. The starved *T. castaneum* adults show in a series of the behavior from recognition, eating to excretion the gypsum diet. Scale bars: 1 mm. **A.** The adults bit gypsum. The gypsum was stained with Coomassie Brilliant Blue (CBB). **B, B’.** Digestive tract of adult beetle with the CBB staining gypsum **(B)** and without the CBB staining **(B’). C.** Excreta of the CBB staining gypsum by *T. castaneum* (arrowheads). **D.** The effect of gypsum intake with sugar or sugar alcohol mixture. Each sugar/sugar alcohol at 200 mM was contained in the gypsum block. *T. castaneum* adults fed on the artificial diet for 48 h. The amount of excretion was measured using microbalance. Each plot represents the amount of excreta of adult beetles in individuals (n = 4-10). Standard error bars show S.E.M. Statistical analyses were performed one-way ANOVA and the post hoc Tukey’s multiple comparison test (“**” P<0.0l, “***” P<0.001;, “****” P<0.0001”, “ns” no significant). Fructose, Fru; glucose, Glu; galactose, Gal; sucrose, Suc; maltose, Mal; trehalose, Tre; mannitol, Man; sorbitol, Sor.

### Phylogenetic analysis of insect Grs

Mannitol, a sugar alcohol, acts as a sweetener for vertebrate chemosensation [24], and other sugar alcohols are readily recognized by insect species. In *B. mori*, sugar alcohols such as *myo/epi*-inositol were recognized by the gustatory receptor BmGrlO belonging to the Gr43-like clade [9]. We aimed to identify homologous genes in *T. castaneum* belonging to the same clade as BmGr10 based on the phylogenetic analyses using amino acid residues. The phylogenetic tree of the insect Grs was constructed using neighbor-joining (Fig 2). TcGr20, 21, and 25-28 belonged to the clade including the DmGr43a fructose receptor derived from *Drosophila melanogaster*, the BmGr9 fructose receptor and BmGr10 *myo/epi-*inositol receptors derived from *B. mori*, and the HarmGR4 fructose receptor derived from *Helicoverpa armigera* [9, 11, 20]. On the basis of these results, we hypothesized that mannitol receptors belonged to the TcGr20, 21, and 25-28 groups.

**Fig 2.**
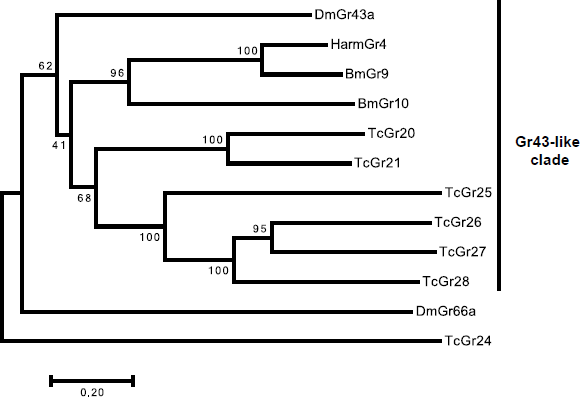
Phylogenetic analysis of deduced amino acid sequences of Gr43-like genes. Amino acid sequence alignment was generated using ClustalW, and a rooted tree of insect Grs was conducted by neighbor-joining method in MEGA ver. 7 [43]. The percentage of replicate trees are shown in the associated taxa clustered together in the bootstrap test of 500 replicates. The scale bar represents 0.2 substitutions per amino acid site. Amino acid sequences of *B. mori* and *T. castaneum* Grs were obtained from [14, 44], respectively. The other amino acid sequences, AmGr3 derived from *Apis mellifera*, HarmGr4 from *Helicoverpa armigera*, DmGr43 and DmGr66a derived from *Drosophila melanogaster*, were obtained from the NCBI public database.

### TcGr20 is a mannitol/sorbitol receptor

We attempted to amplify the ORF in the *TcGr20, 21*, and *25-28* genes for subcloning into the *Xenopus* oocyte expression vector. However, we failed to obtain PCR products of *TcGr25* and *26* from any cDNA produced from any sampled tissues and stages. Hence, we analyzed the biochemical functions of TcGr20, 21, 27 and 28 using electrophysiological analysis. Two-electrode voltage-clamp recording was performed in accordance with a previous study [9]. We reconfirmed a BmGr10 response to myo-inositol using a two-electrode voltage clamp as a positive control (S1 Fig). Each *TcGr20,21,27* and 28-expressing oocyte was clamped with electrode capillaries filled with 3 M KCl. When mannitol was added to the perfusion chamber, an inward current was observed in *TcGr20-ex*pressing oocytes (Fig 3A), but not in *TcGr21-, TcGr27-* or *TcGr28-ex*pressing or water-injected oocytes (Fig 3B, S2 Fig). TcGr20 also responded to sorbitol but not the other sugars (Fig 3A). We did not observe a current response in *TcGr21, 27* or 28-expressing oocytes for any sugars/sugar alcohols tested. The mannitol/sorbitol-induced currents observed in TcGr20 expressing oocyte were concentration-dependent to 20-200 mM mannitol and 40-230 mM sorbitol (Fig 4A, 4B). Based on the dose-response curves, the EC50 value of mannitol and sorbitol were 72.6 ± 9.1 mM and 90.6 ± 10.4 mM, respectively (Fig 4C, 4D). This result suggests that TcGr20 functions as a mannitol/sorbitol receptor.

**Fig 3.**
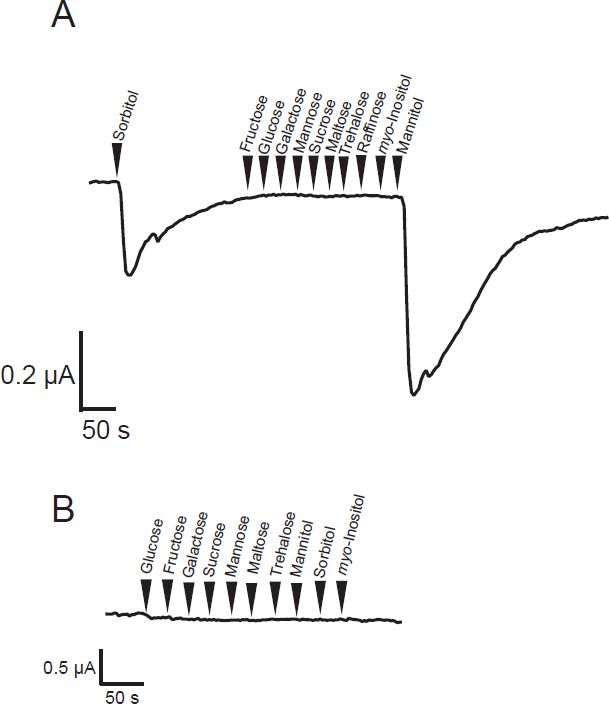
Current recording of *Xenopus* oocytes expressing TcGr20. **A.** Inward current response of *Xenopus* oocytes expressing TcGr20 to candidate tastants (arrowheads). Tastants were tested at 200 mM. **B.** The current of water-injected oocytes to same tastants were also recorded. The current data are representative of recordings independently performed in several times.

**Fig 4.**
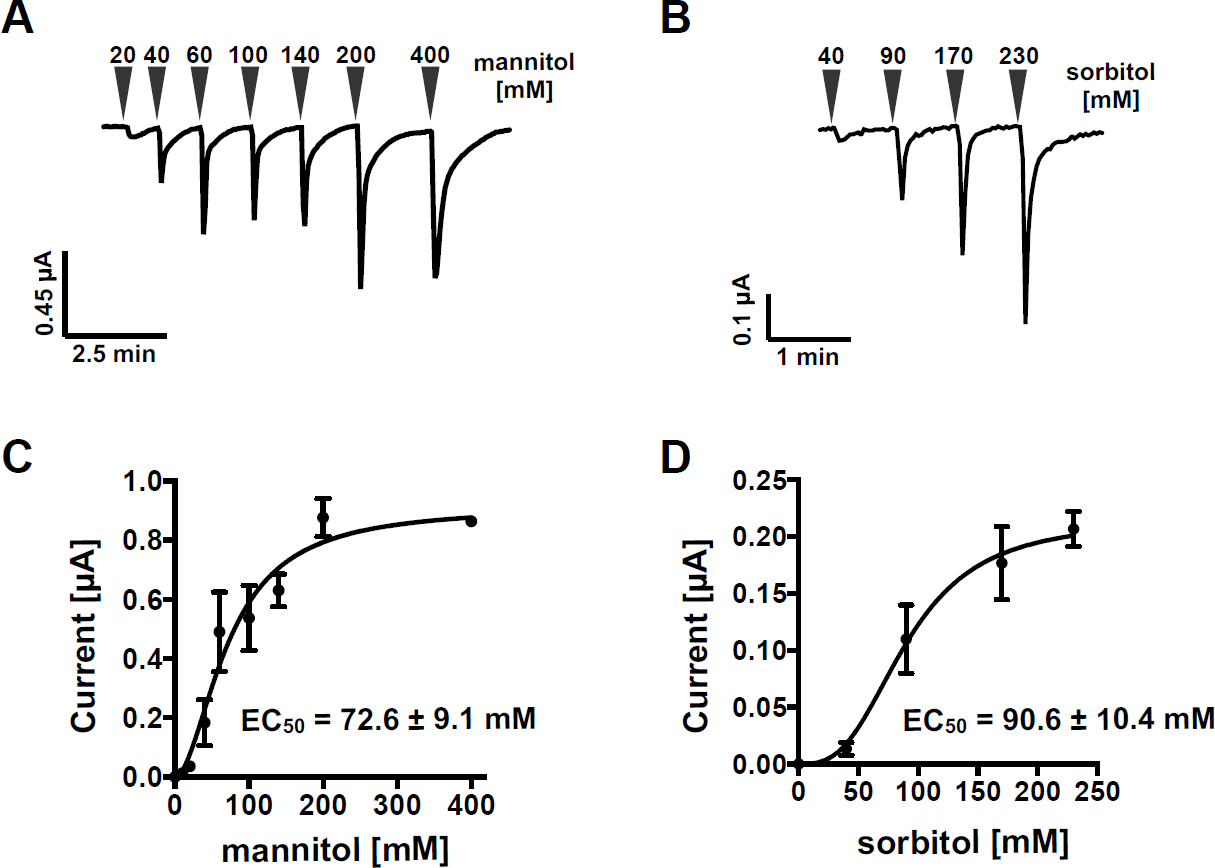
Ligand dose-dependent response ofTcGr20. Two-electrode voltage clamp recordings of TcGr20-expressing *Xenopus* oocytes. **A-B.** Inward current response of TcGr20-expressing oocytes with a range of 20 to 400 mM mannitol **(A),** and 40 to 230 mM sorbitol **(B).** Each arrowhead represents various concentrations. **C-D.**Curves were fitted with a standard slope, and EC values were calculated for mannitol **(C)** and sorbitol **(D),** respectively. Data are shown as mean ± S.E.M. (n = 3).

### Tissue expression of *TcGr20*

Previous studies have shown that *TcGr20* is predominantly expressed in the antennae with RNA sequencing and *in situ* RT-PCR [15, 25]. Our findings here confirm that *TcGr20* is mainly expressed in the antennae (Fig 5), as well as in head structures including the mouthparts, and in the thorax, abdomen, and legs in both males and females. *TcGr20* was highly expressed in antennae in both male and female adults (Fig 5A, 5B). TcGr20 on the antennae likely acts as sensors for mannitol and sorbitol.

**Fig 5.**
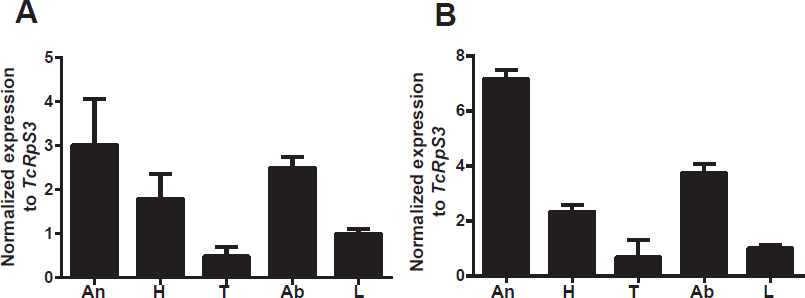
Tissue expression of *TcGr20*. The relative expression level of the *TcGr20* in different tissues was determined by quantitative RT-PCR. *TcGr20* expression in male **(A)** and female **(B).** An, antennae; H, head; T, thorax; Ab, abdomen and L, legs. Relative expression was caluclated using *ΔΔCt* method. Ribosomal protein S3 (*RpS3*) in *T. castaneum* was used as the control to normalize the amount of templates. Data are shown as mean ± S.E.M. (n = 3).

### Effect of mannitol/sorbitol concentrations in the dietary intake assay

We measured the dietary intake of *T. castaneum* adult beetles using gypsum containing mannitol or sorbitol at various concentrations (Fig 6). The dietary intake increased in the presence of 100 mM mannitol (Fig 6A) and 200 mM sorbitol (Fig 6B), respectively. The amount of gypsum excreta in individuals at 48 h was 0.6 ± 0.07 mg for 100 mM mannitol, and 0.26 ± 0.03 mg for 200 mM sorbitol (Fig 6). These results indicate that mannitol stimulates a feeding response even at lower concentrations, promoting dietary intake by the beetles.

**Fig 6.**
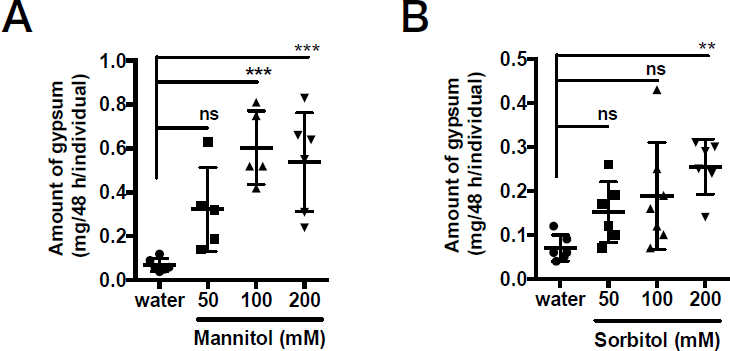
Concentration response of mannitol and sorbitol in TribUTE assay. The dose effect of gypsum intake with sugar alcohol mixture. A, mannitol; B. sorbitol. Sugar alcohols at various concentrations were contained in the gypsum block. *T. castaneum* adults fed on the gypsum for 48 h. The amount of excreta was measured using microbalance. Scatter plot represents the amount of excretion of adult beetles in individuals (n = 5-7). Standard error bars show S.E.M. Statistical analyses were performed one-way ANOVA and the post hoc Tukey’s multiple comparison tests (“**” P<0.01, “***” P<0.001, “ns” no significant).

### Evaluation of dietary intake in *TcGr2*0-silencing *T. castaneum*

Our results showed that *T. castaneum* fed on gypsum in the presence of mannitol (Fig 6A), and that TcGr20 was a mannitol receptor (Fig 3A). We therefore next investigated whether TcGr20 was involved in mannitol recognition for feeding. One promising approach for validating gene function, RNA interference (RNAi), was effective in *T. castaneum* [26]. We injected *TcGr20* double-strand RNA (dsRNA) into *T. castaneum* adults; as a result, *TcGr20* expression was significantly suppressed compared with that of *emerald luciferase (Eluc)-*dsRNA injected adults (Fig 7A). Using the *TcGr20* dsRNA-injected adult beetles, we evaluated the dietary intake of gypsum in the presence of 100 mM mannitol. The amounts of excreta from *TcGr20* dsRNA-injected adults significantly decreased in the presence of 100 mM mannitol compared in comparison to that of *Eluc* dsRNA-injected adults (Fig 7B). These results indicate that *TcGr20* RNAi is an effective tool, and that TcGr20 is responsible for mannitol recognition *in vivo*.

**Fig 7.**
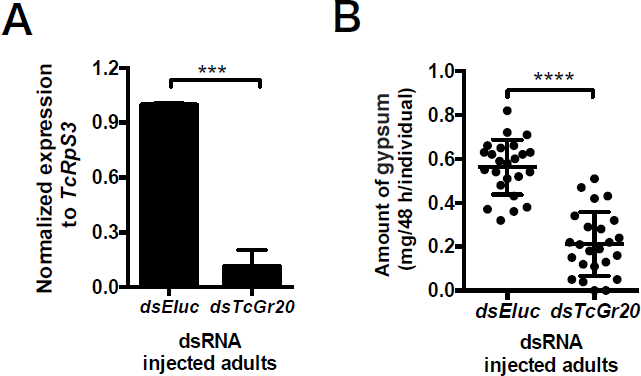
TribUTE assay in *TcGr20*-silencing *T. castaneum*. **A.** Knockdown of *TcGr20* by using the injection of the dsRNA into the starved adult beetles. The *Eluc-*dsRNA was injected as a control. *TcGr20* expression levels of whole body were examined at 48 h after the dsRNA injection. Standard error bars show S.E.M. Statistical significance was determined by *t*-test (P = 0.0006). **B.** Effect of gypsum intake in the presence of 100 mM mannitol. The gypsum in the presence of 100 mM mannitol was given to the *TcGr20-* and *Eluc-*dsRNA-injected adult beetle individuals at 48 h after dsRNA injection, respectively for 48 h. Amount of gypsum was measured as excretes. Each plot represents the amount of excreta of adult beetles in individuals (n = 24). Standard error bars show S.E.M. Statistical significance was determined by *t*-test (“****”P < 0.0001).

## Discussion

Our results demonstrate that mannitol acts as a significant feeding attractant in *T. castaneum* adult beetles, and that TcGr20 is responsible for mannitol/sorbitol recognition and the promotion of dietary intake.

Gustatory receptor (Gr)-expressing chemosensory organs are involved in external non-volatile compound recognitions. Antennae and legs recognize sucrose in *Tribolium brevicornis* in electrophysiological analyses [27]. In *H. armigera*, the fructose receptor HarmGr4 in the antenna recognized fructose [11]. Hence, *TcGr* genes expressed in antennae or legs were hypothesized as being involved in the perception of mannitol as an external signal. In the present study, we used qRT-PCR analysis to show that *TcGr20* was expressed in the antennae. Previous analyses such as tissue-specific RNA-seq and *in situ* PCR have also shown that *TcGr20* is expressed in the antennae [15, 25], and our results are consistent with these findings. We observed a difference in *Gr* expression levels between males and females [28], implying the occurrence of sex-specific feeding behavior. However, we observed no difference in *TcGr20* expressions between the sexes [Fig 5).

Electrophysiological analyses using *Xenopus* oocytes showed that TcGr20 contributes to responses to mannitol and sorbitol (Fig 3). The EC_50_ values of TcGr20 for these sugar alcohols showed that the mannitol response was 0.8 times more sensitive than the sorbitol response. Additionally, the dietary intake of *T. castaneum* adult beetles in the TribUTE assay was also more sensitive to mannitol than sorbitol (Fig 6). Mannitol response was significant at 100 mM, while the significance of sorbitol response was observed at 200 mM (Fig 6), indicating that the concentration response of mannitol appears to be correlated with that of TcGr20 response levels in the electrophysiological analysis. It is likely that TcGr20 mainly regulates mannitol recognition in the gustatory organs. We also examined whether TcGr20 is involved in mannitol recognition using RNAi and the TribUTE assay. To explore effective RNAi in *T. castaneum, TcGr20-dsRNA* was injected into pupae; however, the *TcGr20* expression levels in *TcGr20*-dsRNA-injected adult beetles after eclosion showed a negligible difference compared to those injected with *Eluc*-dsRNA injection (S3 Fig). The 6-9-day period of pupal development prior to adulthood [29] may have decreased the gene silencing effect To test this possibility, we attempted dsRNA injection into starved adult beetles. The test showed a significant RNAi silencing effect (Fig 7A). Using the *TcGr20* RNAi beetles (Fig 7A), we showed that TcGr20 is responsible for mannitol recognition-dependent dietary intake behavior (Fig 7B). The attraction of *TcGr20*-dsRNA-injected beetles to the gypsum block appeared to decrease in the presence of 100 mM mannitol, resulting in a decrease in gypsum intake. It is important to note that the silencing effects of RNAi were obtained temporarily. TcGr20 responds to both mannitol and sorbitol, but these dietary intake assays should be further confirmed using other methods such as the clustered regularly interspaced short palindromic repeats (CRISPR)/Cas9 system.

In *T. castaneum*, six *TcGr* genes in the *Tribolium* genome (NCBI public data, ID: 216) were found as the Gr43-like clade (Fig 2). This study demonstrated that TcGr20 is a mannitol/sorbitol receptor, but the functions of other *TcGr21-28* genes remain unclear. These include *TcGr25* and *26*, for which we failed to amplify PCR products from any cDNAs. The six *TcGr* genes would allow *T. castaneum*, as a generalist feeder, to be capable of recognizing a large number of natural attractants such fructose and sugar alcohols. Since these *TcGr* genes were highly expressed in the antennae, contributing to the beetles’ ground-dwelling life style and scanning behavior [15], they may respond to non-volatile stimuli produced by host plants. As alternative candidate targets, we expected that compounds produced by fungi would be attractive, since beetles were strongly attracted to chemical stimuli from fungi grown on flour and cotton seeds [30-33]. When the beetle larvae fed on *Aspergillus niger*, suitable development occurred and the reproductive potential of the females eventually increased (30-32).

Mannitol, the most widely distributed of the polyols, is found in more than 50 species of plants, algae, fungi, and lichens [34, 35]. In particular, dried seaweeds contain 1-1.7 M mannitol as a major carbohydrate component [36]. Sorbitol is also present in prunes at 2.4 g/100 g dry weight and pears at 4.6 g/100 g dry weight [37]. *T. castaneum* would recognize these stored foods because sugar alcohols are present at much higher concentrations than are responded to by TcGr20. *A. niger* produces 45-210 mM mannitol [38], implying that *T. castaneum* can recognize the fungi themselves using TcGr20. Fungal growth is favored in high-moisture conditions in stored products and bulk grains. It is possible *T. castaneum* may be drawn to such stored products using such fungal attractants as a cue. *T. castaneum* is also a harmful wheat flour pest [12, 39], attacking cake flour and whole-wheat flour containing around 0.032 mg/g and 0.01 mg/g dry weight mannitol/flour, respectively (S4 Table), which are levels far lower than biochemically required for TcGr20 recognition. Therefore, *T. castaneum* adults are unlikely to recognize wheat flour using TcGr20. Rather, they more likely recognize sugars such as glucose and fructose contained in the wheat flour. Our TribUTE assay demonstrated that adult beetles could, to some degree, recognize some sugars such as fructose, glucose, maltose, and trehalose (Fig 1D). Based on these findings, we suggest that TcGrs in the sugar clade [15] are potential candidates for recognition of these carbohydrates contained in wheat flour.

A novel artificial dietary method for *T. castaneum* adult beetles facilitates examination of their dietary intake based on the eventual measurement of amount of excreta. Gypsum, or calcium sulfate dihydrate, is an inorganic compound, formed into a solid in combination with water (PubChem database, CID: 24928). The observation of gypsum intake by *T. castaneum* demonstrated a series of behaviors from recognition and swallowing to excretion. The TribUTE assay can quantify beetle preferences for non-volatile compounds by measuring gypsum excreta without consideration of odorant stimulation. Sugars/sugar alcohols act as feeding behavior-facilitating factors in some insects [40-42], and *T. castaneum* significantly preferred gypsum containing these additives (Fig 1D). The TribUTE assay enabled the identification of non-volatile compounds associated with the food preference of *T. castaneum*. Notably, adult beetles can feed on gypsum without sweeteners (Fig 1). Thus, the TribUTE assay would be also applicable for the exploring non-volatile compounds that induce a decrease in dietary intake by *T. castaneum*. Such compounds would be useful in controlling *T. castaneum* in the future.

## Author contributions

TT, SK conceived and designed the experiments; TT, SK performed the experiments; TT, RS, SK analyzed data; TT SK contributed reagents/materials/analysis tools; TT, SK wrote the paper.

## Conflicts of interest

The authors declare no competing financial interests.

## Acknowledgements

We would like to thank Dr. Takumi Kayukawa at Institute of Agrobiological Sciences NARO (Ibaraki, Japan) for advice of *T. castaneum* maintenance. We appreciate to Dr. Takeshi Suzuki at Tokyo University of Agriculture and Technology for the quantifications of DNA and RNA.

## Supporting information

S1 Table. Primers for *Xenopus* oocyte expression

S2 Table. Primers for quantitative RT-PCR

S3 Table. Primers for double strand RNA synthesis

S4 Table. Mannitol in flours

S1 Fig. The BmGr10 response to myo-inositol as a positive control in a two-electrode voltage clamp

S2 Fig. Current recordings of *Xenopus* oocytes expressing TcGr21, TcGr27 and TcGr28 against sugars and sugar alcohols

S3 Fig. Gene silencing effect

S1 file. Quantification of mannitol content in wheat flours

